# Evidence for shared ancestry between Actinobacteria and Firmicutes bacteriophages

**DOI:** 10.1101/842583

**Authors:** Matthew Koert, Júlia López-Pérez, Courtney Mattson, Steven Caruso, Ivan Erill

## Abstract

Bacteriophages typically infect a small set of related bacterial strains. The transfer of bacteriophages between more distant clades of bacteria has often been postulated, but remains mostly unaddressed. In this work we leverage the sequencing of a novel cluster of phages infecting *Streptomyces* bacteria and the availability of large numbers of complete phage genomes in public repositories to address this question. Using phylogenetic and comparative genomics methods, we show that several clusters of Actinobacteria-infecting phages are more closely related between them, and with a small group of Firmicutes phages, than with any other actinobacteriophage lineage. These data indicate that this heterogeneous group of phages shares a common ancestor with well-defined genome structure. Analysis of genomic %GC content and codon usage bias shows that these actinobacteriophages are poorly adapted to their Actinobacteria hosts, suggesting that this phage lineage could have originated in an ancestor of the Firmicutes, adapted to the low %GC content members of this phylum, and later migrated to the Actinobacteria, or that selective pressure for enhanced translational throughput is significantly lower for phages infecting Actinobacteria hosts.

## Introduction

Frequently referred to as phages, bacteriophages are viruses capable of infecting bacteria. It has been estimated that phages are the most abundant entities in the biosphere (Fokine and Rossmann 2014) and, through their regulation of bacterial populations, bacteriophages play an essential role in many global processes of the biosphere, such as carbon and nitrogen cycling (Casjens 2005). In the last decade, decreasing sequencing costs have dramatically increased the number and diversity of bacteriophage genome sequences (Russell and Hatfull 2017). This influx of phage genomic data has reinforced the notion that phages are not only key players in geobiological processes, but also the largest reservoirs of genetic diversity in the biosphere (Pedulla et al. 2003). The Science Education Alliance-Phage Hunters Advancing Genomics and Evolutionary Science (SEA-PHAGES) program has undertaken a sustained effort to isolate and sequence phages infecting Actinobacteria species (Russell and Hatfull 2017). Among these, mycobacteria-infecting phages have been studied the most, providing a remarkably deep sample of bacteriophages infecting a given bacterial genus (Russell and Hatfull 2017). Studies of genetic diversity in over 600 mycobacteria-infecting phage genomes have revealed extensive mosaicism, and genetic exchange among relatively distant groups of mycobacteriophages. Rarefaction analyses suggest that the mycobacteriophage gene pool is not an isolated environment, and that it is enriched by an influx of genetic material from outside sources (Pope et al. 2015). Here we report on the genomic characterization of a new cluster of *Streptomyces* phages (Cluster BI). Gene content and protein sequence phylogenies indicate that members of BI and related actinobacteriophage clusters share a common ancestor with *Lactococcus* and *Faecalibacterium* phages (Garneau, Tremblay, and Moineau 2008; Kot et al. 2014). Analysis of genomic %GC content and codon usage bias indicates that these actinobacteriophages are still undergoing amelioration, suggesting that selective pressure for translational optimization is weak, or that they could have originated as a result of an interphylum migration event from related Firmicutes phages.

## Methods

### Genome data

Genomes for relevant *Streptomyces* phages and for reference Actinobacteria and Firmicutes bacteriophages were retrieved in GenBank format from the NCBI GenBank database (Benson et al. 2017) using custom Python scripts. These scripts also derived nucleotide and amino acid FASTA-formatted files from the GenBank records, and autonumerically reassigned locus_tag and gene GenBank identifiers for consistent pham annotation with PhamDB. For phages without a public GenBank record, nucleotide FASTA files were downloaded from PhagesDB (Russell and Hatfull 2017) and auto-annotated with DNA Master (Pope and Jacobs-Sera 2018) to generate a GenBank-formatted file. For %GC analysis and CUB analyses, host reference genomes were obtained at the strain, species or genus level, based on availability. Cluster assignments for Actinobacteria-infecting phages were obtained from PhagesDB (https://phagesdb.org/), which systematically classifies database phages into clusters according to the fraction of shared proteome (>35%) (Pope et al. 2017).

### %GC content and CUB analysis

%GC content data was obtained from the corresponding NCBI assembly records. Group %GC content was compared using a Mann-Whitney U test with α=0.05 using a custom Python script and the scipy.stats module. Codon usage bias was measured using nRCA, a codon adaptation index that compensates for mutational biases and reflects primarily translational selection bias (O’Neill, Or, and Erill 2013). A reference genome was selected for each host bacterial genus and a self-consistent reference set for this host was detected using an expectation-maximization procedure (Data S1) (O’Neill, Or, and Erill 2013). Using these reference sets, for each host and phage genome in a given genus, an nRCA value was obtained for each protein-coding gene sequence, and genome-wide nRCA values were computed as the average across all protein-coding genes.

### Gene content phylogeny

PhamDB was used to compute protein families, or phams, for the bacteriophage genomes under analysis (Lamine, DeJong, and Nelesen 2016). The PhamDB-generated database was then imported into Phamerator (Cresawn et al. 2011) and the resulting pham table was exported as a comma-separated file and processed with spreadsheet software and the Janus program (Lawrence Lab) to obtain a Nexus-format file with presence/absence of each pham in each genome as a binary character. This Nexus file was used as input for SplitTree (Kloepper and Huson 2008). Network and tree phylogenies were inferred with the NeighborNet and BioNJ algorithms using a gene content distance (Snel, Bork, and Huynen 1999) and branch support for the resulting phylogeny was estimated from 1,000 bootstrap pseudoreplicates. A genome-based phylogeny was generated with the VICTOR webservice (Meier-Kolthoff and Göker 2017). Intergenomic protein sequence distances were computed with 100 pseudo-bootstrap replicates using the Genome-BLAST Distance Phylogeny (GBDP) method optimized (distance formula d_6_) for prokaryotic viruses (Meier-Kolthoff and Göker 2017; Meier-Kolthoff et al. 2013) and a minimum evolution tree was computed with FASTME on the resulting intergenomic distances (Lefort, Desper, and Gascuel 2015).

### Protein sequence phylogeny

A profile Hidden Markov Model (HMM) of terminase protein sequences was built with HMMER (hmmbuild) using a ClustalW multiple sequence alignment of all annotated terminase, TerL or terminase large subunit sequences in the genomes under analysis (Eddy 2011; Sievers et al. 2011) (Data S2). This profile HMM was used to search (hmmsearch) the protein FASTA file derived from each genome with a cutoff e-value of 10^−3^. Putative terminase sequences identified by the profile HMM were aligned with ClustalW using default parameters. Tree inference was performed on the resulting multiple sequence alignment using the BioNJ algorithm with a Gamma distribution parameter of 1 and the Jones-Taylor-Thornton substitution model, and branch supports were estimated from 1,000 bootstrap pseudoreplicates (Gascuel 1997). Additional Maximum-Likelihood tree inference was performed with IQ-TREE 2 (Minh et al. 2020), using a General Variable Time matrix model with empirical base frequencies and FreeRate heterogeneity (Müller and Vingron 2000; Le, Dang, and Gascuel 2012), automatically selected by IQ-TREE 2, and 1000 UltraFast bootstrap alignments (Minh, Nguyen, and von Haeseler 2013).

## Results

### Conserved architecture of BI cluster *Streptomyces* phage genomes

In the last few years, our group has characterized and sequenced several *Siphoviridae* bacteriophages capable of infecting *Streptomyces scabiei* RL-34 (Blocker et al. 2018). Genomic analysis indicated that these bacteriophages belong to the PhagesDB BI cluster, which also encompasses bacteriophages isolated by other teams on different *Streptomyces* hosts, such as *Streptomyces lividans* JI1326 (*Streptomyces* phage Bing) or *Streptomyces azureus* NRRL B-2655 (*Streptomyces* phage Rima). Cluster BI phages have linear genomes ranging from 43,060 to 57,623 bp, encompassing from 55 to 91 protein coding genes and no predicted tRNA genes. Comparative analysis of these bacteriophage genomes (Figure 1) reveals nucleotide sequence conservation to be predominant only in the virion structure and assembly genes module, which presents a genetic arrangement consistent with that observed in other *Siphoviridae*, such as PhagesDB cluster J mycobacteriophages (Pope et al. 2013; Lopes et al. 2014). Within this module, the terminase gene shows the highest degree of sequence conservation, followed by segments of the portal, capsid maturation and tape measure protein coding genes (Figure 1). Beyond the structure and assembly module, moderate nucleotide sequence conservation is only observed for the genes coding for a predicted hydrolase in the lysis module, and for the DNA primase/polymerase and a helix-turn-helix (HTH) domain-containing protein in the replication module.

**Figure 1.**
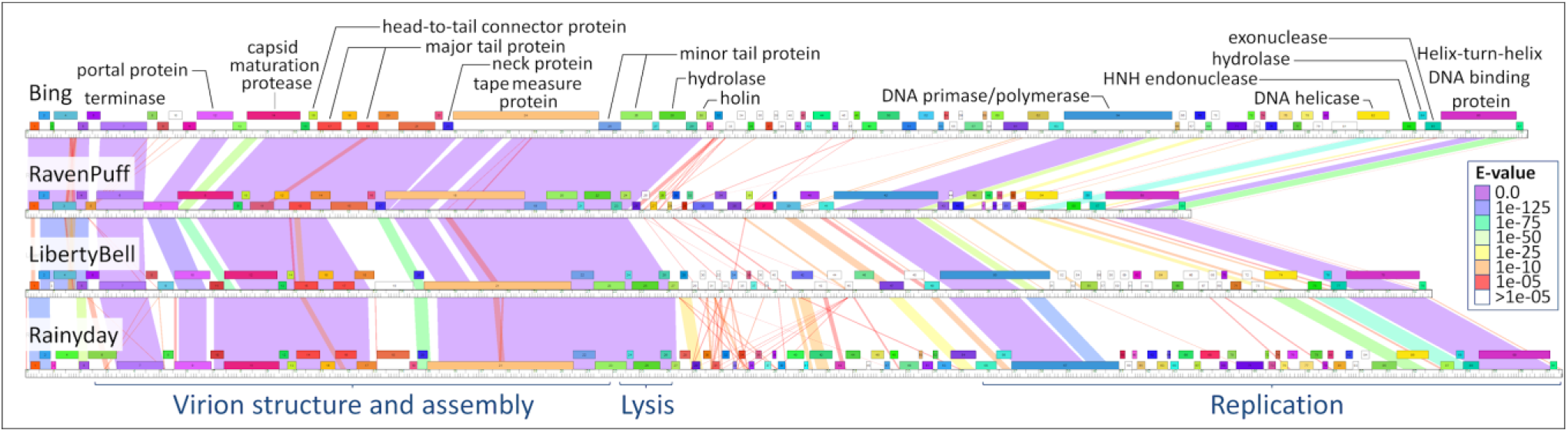
Phamerator-generated map of four representative BI cluster *Streptomyces* phage genomes (Bing (BI1), RavenPuff (BI2), LibertyBell (BI3) and Rainydai (BI4)). Shaded areas between genomes indicate nucleotide similarity, following a purple-to-red rainbow palette that indicates the e-value of the pairwise BLASTn alignment. Genes in the forward strand are shown as boxes above the genome position ruler for each phage; genes in the reverse strand are shown below the ruler. Groups of orthologous protein sequences are denoted by arbitrarily colors in protein-coding gene boxes. Orphams (proteins in a pham containing a single member) are shown as white boxes.

### Interphylum conservation of structure and replication proteins

Functional annotation of BI cluster genomes was performed using BLASTP searches against both the NCBI GenBank and the PhamDB databases, as well as the HHpred service (Söding, Biegert, and Lupas 2005; Russell and Hatfull 2017; Benson et al. 2017; Blocker et al. 2018). During the annotation process, BI cluster protein sequences frequently elicited significant hits against *Arthrobacter* (clusters AM, AU and AW), *Gordonia* (cluster DJ), *Rhodococcus* (cluster CC) and *Microbacterium* (cluster EL) bacteriophages, rather than against other *Streptomyces* phage clusters. It was also noticed that BLASTP searches against NCBI bacterial genomes often returned significant hits against putative prophages in several Firmicutes genomes. This prompted us to search for potential homologs of BI cluster proteins in the genomes of bacteriophages isolated from Firmicutes hosts, and we identified several *Lactococcus lactis* bacteriophage genomes related to *Lactococcus* phage 1706 (Garneau, Tremblay, and Moineau 2008; Kot et al. 2014) and a *Faecalibacterium* phage (FP_oengus, (Cornuault et al. 2018)) harboring multiple homologs of BI cluster proteins.

To contextualize this finding, we compiled complete genome sequences of bacteriophages in all the aforementioned PhagesDB clusters and in the *Lactococcus* and *Faecalibacterium* group, as well as reference members from other *Streptomyces, Arthrobacter, Gordonia* and *Rhodococcus* clusters, reference Firmicutes phages (e.g. *Staphylococcus* virus Twort, *Bacillus* virus SPO1, *Lactococcus* phage P335, *Leuconostoc* phage 1-A4, *Bacillus* phage Bam35c) and other bacteriophages identified by BLASTP as containing proteins with significant similarity to BI cluster proteins. Using PhamDB and Phamerator, we generated a table of orthologous protein sequence groups (phams) across this heterogeneous set of bacteriophage genome sequences (Table S1). A quick assessment of predicted phams revealed that the phams with the largest number of members within this dataset clearly outlined a supercluster of actinobacteriophages encompassing *Arthrobacter* (clusters AM, AU and AW), *Gordonia* (cluster DJ), *Rhodococcus* (cluster CC), *Microbacterium* (cluster EL) and *Streptomyces* (cluster BI) phages. Importantly, 10 out of the 11 phams that are present in all these 41 actinobacteriophages were also found in the *Lactococcus* and *Faecalibacterium* group. Overall, the actinobacteriophage supercluster shared 27 large phams (27.6 ±_SD_ 17.8 members) with the *Lactococcus* and *Faecalibacterium* phage group, and 9 of the 15 largest phams were shared between both groups (Table S1). In contrast, *Lactococcus* and *Faecalibacterium* phages did not present any shared phams with the putatively related *Lactococcus* phage P335 (Garneau, Tremblay, and Moineau 2008), and they only shared five small phams (2 members) with reference Firmicutes phages. Likewise, the identified actinobacteriophage supercluster only shared 35 small phams (6.0 ±_SD_ 3.1 members) with other actinobacteriophages.

Graphical analysis of the genomic distribution of orthologs spanning both the actinobacteriophage supercluster and the *Lactococcus* and *Faecalibacterium* phages (Figure 2) revealed that most of the orthologous genes were contained within two conserved regions at opposite ends of the genome. The first conserved region encompasses a sizable fraction of the virion structure and assembly genes module seen in BI cluster phages, containing a HNH endonuclease, a head-to-tail connector, the terminase large subunit, the portal protein and a capsid maturation protease (Figure 2A). The second conserved region corresponds to the end of the replication module observed in cluster BI phages and contains a DNA helicase, a HNH endonuclease, a RecB exonuclease, the HTH domain-containing protein and a conserved hypothetical protein (Figure 2B). Pairwise amino acid identity and alignment coverage for conserved orthologs among actinobacteriophages were moderately high (56% ±_SD_ 12 and 91% ±_SD_ 7), and remained surprisingly high between *Gordonia* phage Gravy and *Faecalibacterium* phage FP_oengus (49% ±_SD_ 11 and 90% ±_SD_ 7), suggesting a relatively close evolutionary relationship.

**Figure 2.**
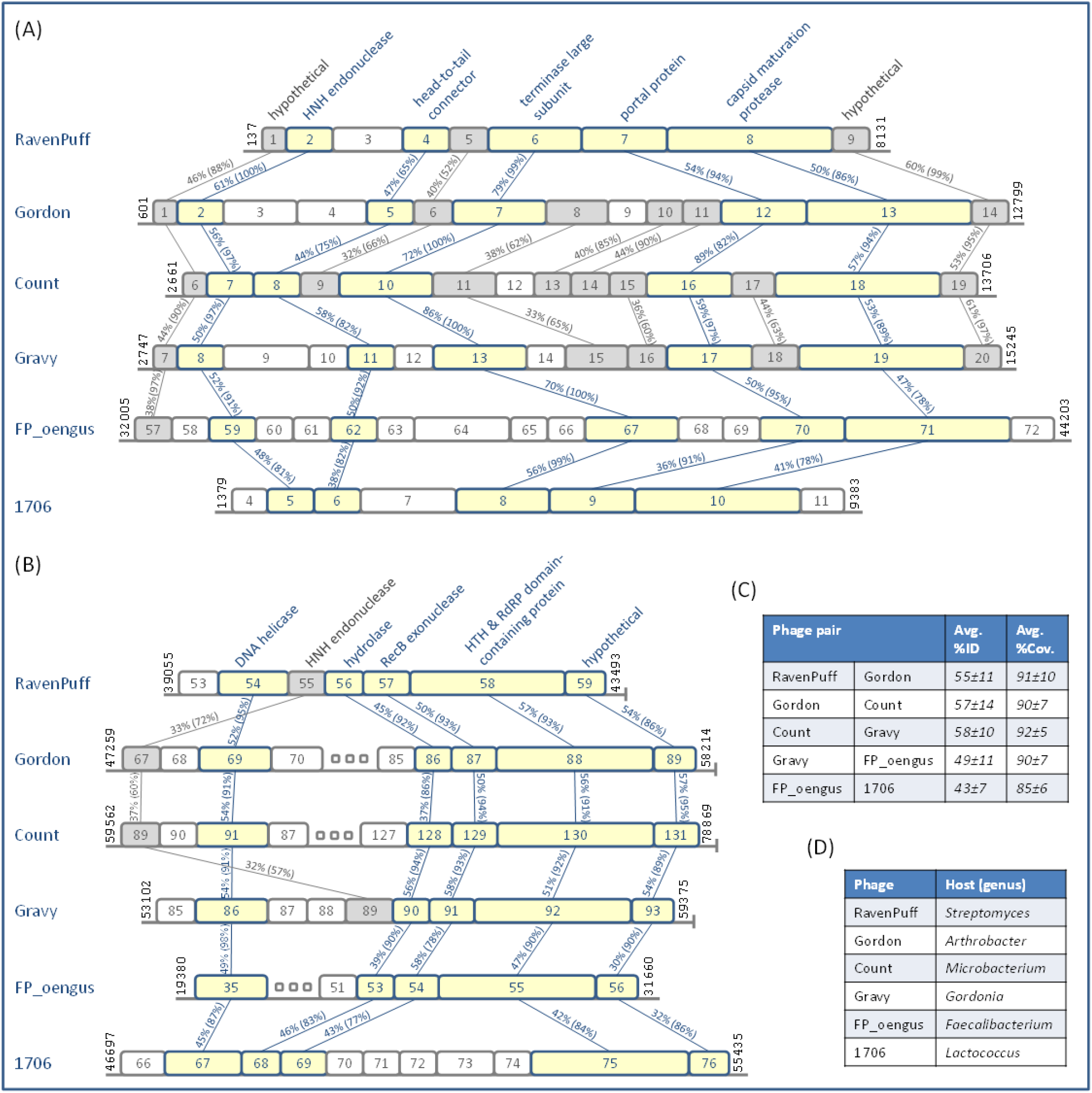
Comparative analysis on representative genomes of the main genomic regions (A and B) containing conserved orthologs. Shaded boxes indicate orthologs conserved in at least two (grey) or in all the species shown (yellow), with the numbers across the lines connecting them showing the pairwise amino acid identity and alignment coverage. Gene numbers and genomic positions are provided for reference in each genome. (C) Average pairwise amino acid similarity and alignment coverage for orthologs conserved acr0ss all species. (D) List of representative phages and their host genera.

The HTH domain-containing protein in the second conserved region (Figure 2B) is annotated in FP_oengus (AUV56548.1) as a putative RNA polymerase. A BLASTP search identified homologs of this sequence only within members of the aforementioned actinobacteriophage supercluster and the *Lactococcus* and *Faecalibacterium* phage group. An HHpred search with their multiple sequence alignment revealed a significant hit (P=95.68%, 286 aligned columns) with the PFAM model PF05183.13 (RdRP; RNA-dependent RNA polymerase), as well as the presence of HTH-based DNA binding domains at both the N- and C-terminal ends, which was confirmed with two HTH prediction services (Dodd and Egan 1990; Narasimhan et al. 2002). Close examination of the multiple sequence alignment revealed the presence of two RNA-polymerase sequence motifs described recently for crAss-like family phages and YonO-like RdRP homologs (Iyer, Koonin, and Aravind 2003; Yutin et al. 2018), including the signature catalytic loop motif DxDGD shared by RDRPs and DNA-dependent RNA polymerases (Figure S1). This protein could therefore potentially have RNA polymerase activity and hence represent a signature genetic element of this heterogeneous group of phages.

### Shared ancestry between Actinobacteria and Firmicutes phages

The presence of two genomic regions showing substantial numbers of orthologous genes across a group of actinobacteriophages infecting multiple hosts and a small set of Firmicutes phages strongly pointed to an evolutionary relationship among these phages. To validate and examine this hypothesis, we used SplitsTree to infer the neighbor tree and estimate bootstrap support for the splits. The results (Figure 3, Figure S2, Data S3) show consistent branching (99.9% bootstrap support) of the actinobacteriophage supercluster with both *Lactococcus* and *Faecalibacterium* phages, clearly establishing that these Firmicutes phages and the actinobacteriophage supercluster phages share more gene content with each other than with reference actinobacteriophages and Firmicutes phages. To further validate and support this result, we performed phylogenetic inference on the protein sequence of the large terminase subunit (Figure 4, Data S2), a very common marker for bacteriophage phylogenetic analysis (Casjens et al. 2005, 18; Bardina et al. 2016; Merrill et al. 2016; Sharaf et al. 2018). Due to the high diversity among the phages included in the analysis, the alignment of TerL sequences yielded no conserved blocks with GBlocks for Bayesian inference analysis (Castresana 2000). The inferred ML tree therefore provides primarily support for coherent groups of phage sequences. The tree in Figure 4 shows solid support (100% bootstrap support) for a joint branching of the actinobacteriophage supercluster phages and *Lactococcus* and *Faecalibacterium* phages, giving further credence to the notion that these phages share a common ancestor. Identical support for the joint branching of the actinobacteriophage supercluster phages and *Lactococcus* and *Faecalibacterium* phages was obtained through Neighbor Joining tree inference on the TerL alignment (Figure S3; Data S4), and through independent phylogenetic inference using a bootstrapped minimal evolution algorithm operating on intergenomic protein sequence distances inferred from pairwise genome-wide reciprocal tBLASTX (Figure S4).

**Figure 3.**
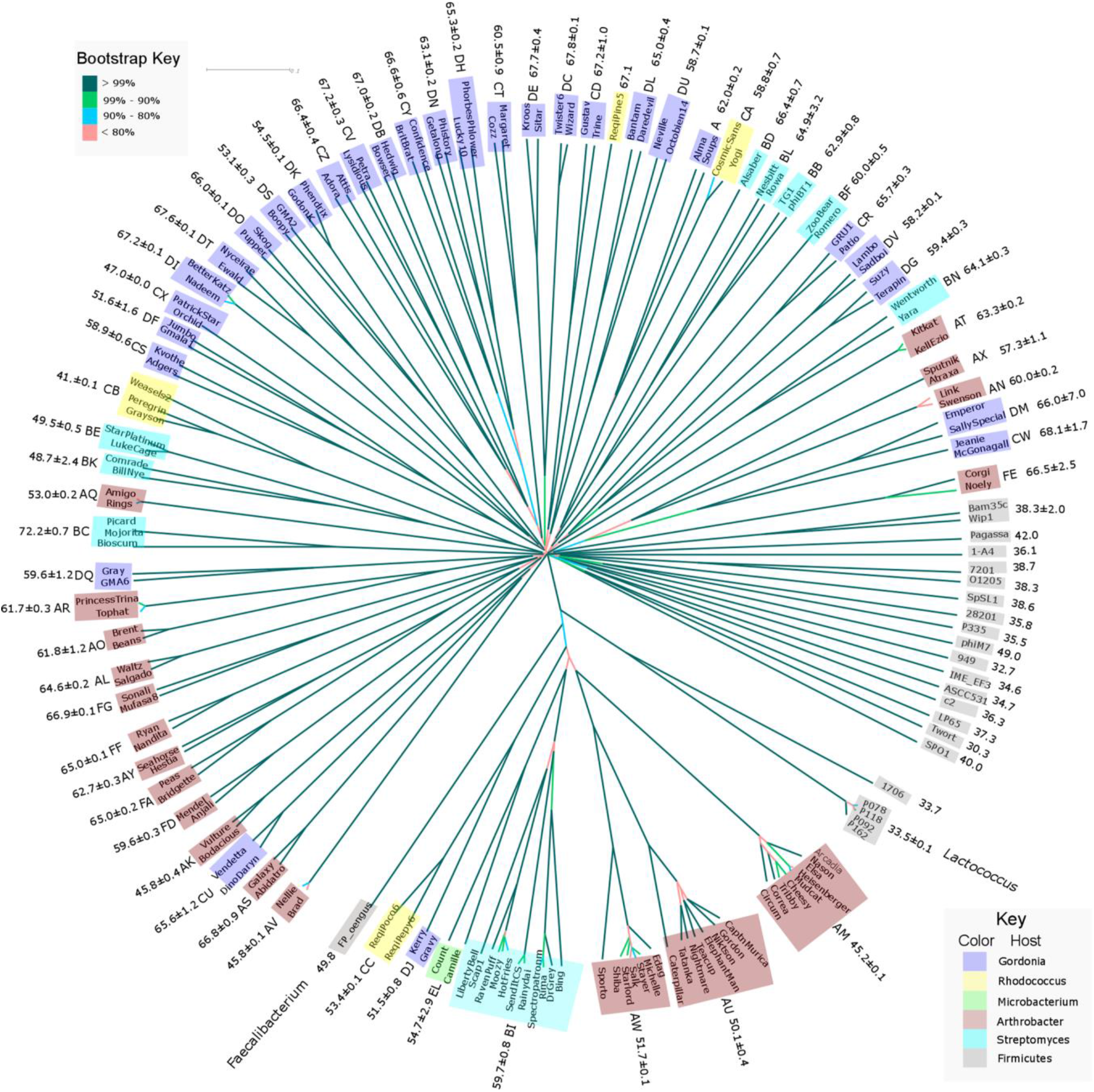
SplitsTree-inferred BioNJ tree for analyzed phages. Bootstrap branch supports for 1,000 pseudoreplicates are shown as percent values on branches. Average genomic %GC content values are shown for different phage groups. Where available, cluster names are also indicated. The phages and pham table used in the analysis are available in Table S1 and Table S2. The Nexus-formatted tree file is available in Data S3.

**Figure 4.**
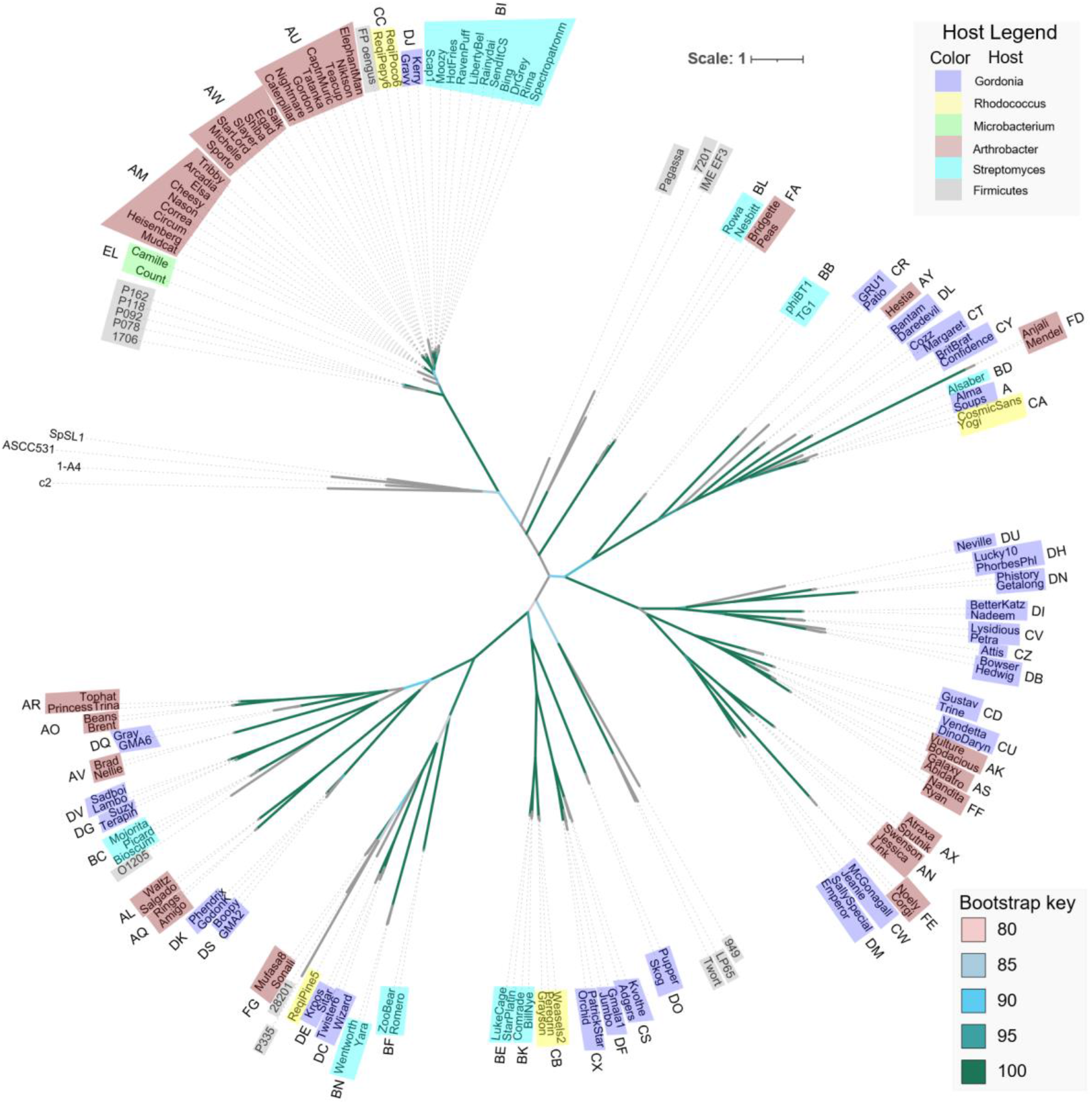
Maximum-likelihood tree for the large terminase subunit protein sequences. Bootstrap branch supports for 1,000 pseudoreplicates are shown as percent values on branches. The phages and terminase sequences used in the analysis are available in Table S1 and Data S2.

The consistent and well-supported branching of actinobacteriophage supercluster phages and *Lactococcus* and *Faecalibacterium* group phages was also confirmed by inspection of recently published large-scale phage phylogenies. A phylogeny of the Caudovirales based on concatenated protein sequences (Low et al. 2019) provides 99% support for the joint branching of all the phages from these two groups used in the analysis: *Arthrobacter* phage Mudcat, *Rhodococcus* phages ReqiPoco6 and ReqiPepy6, and *Lactococcus* phages P078, P118, P162 and P092. Similarly, a taxonomic analysis with the gene network-based vConTACT v.2.0 (Bin Jang et al. 2019) identifies *Lactococcus* phages P078, P118, P162 and P092, as well as *Rhodococcus* phages ReqiPoco6 and ReqiPepy6, forming well-defined genera, and reveals an average fraction of protein clusters (PCs) shared between the members of these two genera and *Arthrobacter* phage Mudcat and *Lactococcus* phage 1706 of 25.38% ±_SD_ 12.66, compared to an average of 1.62% ±_SD_ 0.77 between any of these phages and the 382 phages showing a significant fraction of shared protein clusters with them (Table S3). Lastly, the ViPTree reference tree for dsDNA phages (Nishimura et al. 2017) also depicts *Lactococcus* phages P078, P118, P162, P092 and 1706, *Rhodococcus* phages ReqiPoco6 and ReqiPepy6, and Arthrobacter phage Mudcat forming a well-supported branch (Figure S5).

### Divergence in %GC content and codon usage bias between bacteriophages and their hosts

We analyzed the %GC content of bacteriophage genomes to assess their alignment with the genomic %GC content of their hosts. The results (Figure 5, Table S4) show that, for each genus, the average %GC content of clusters within the actinobacteriophage supercluster is significantly lower (20-30% lower, p<0.05, Mann-Whitney U test) than that of their hosts and also significantly lower (p<0.05) than the average %GC content of other phage clusters infecting the same genera. This is also true for *Faecalibacterium* phage FP_oengus and *Lactococcus lactis* phages, although the difference in %GC content between phages and hosts is much smaller (~5%, p<0.05), as is the difference between supercluster members and other *Lactococcus* phages (~7%, p<0.05). Besides the members of the here identified supercluster, several other actinobacteriophage clusters from PhagesDB (most notably AV, CX, BK, BE and CB) also present %GC content that is significantly lower than the one observed in their natural hosts and than the average for phage clusters infecting their respective genera.

**Figure 5.**
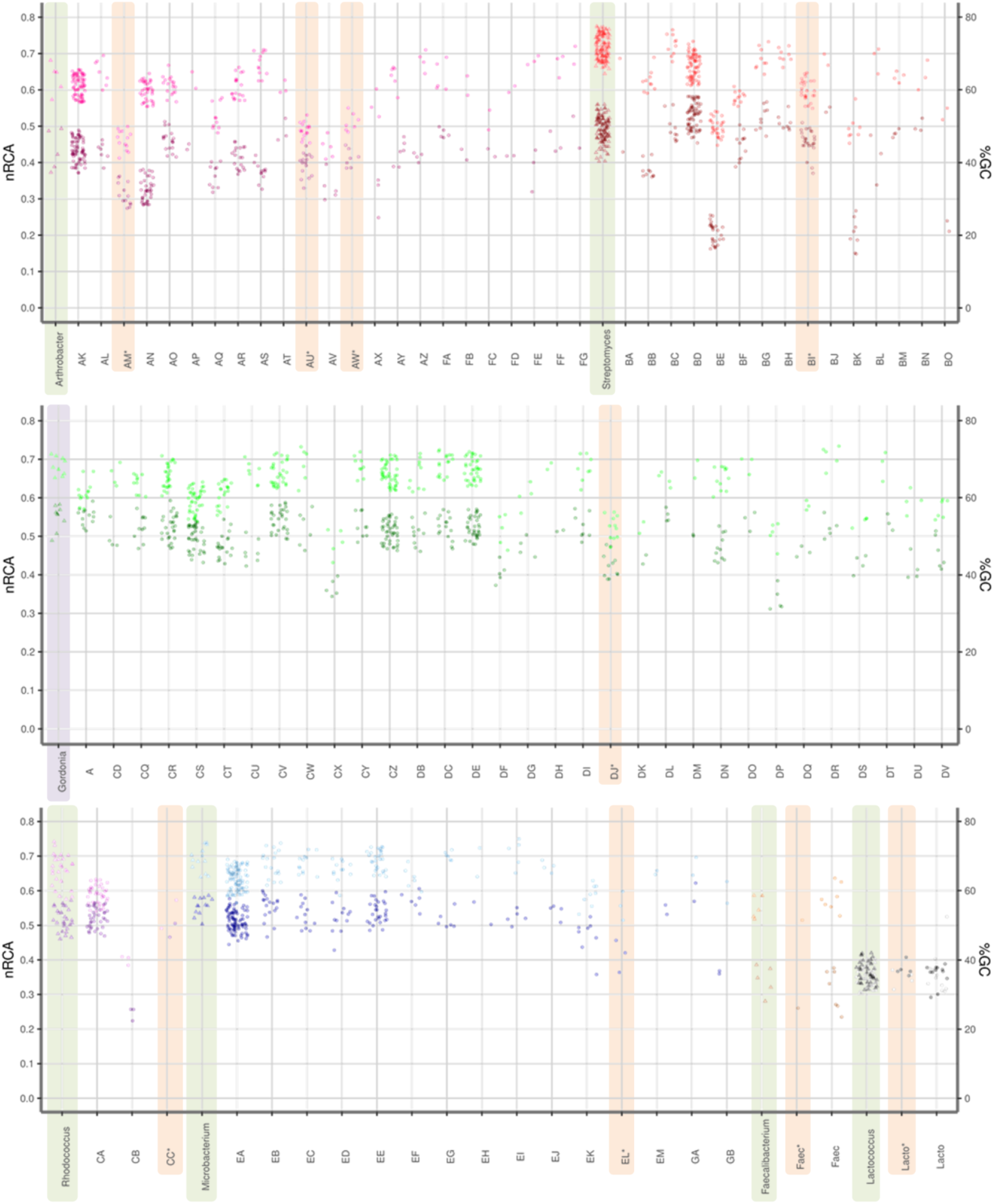
Average %GC and nRCA of phage cluster genomes and of complete genomes from each cluster host genus. Cluster designations for actinobacteriophages are as assigned by PhagesDB. Host data is shown using triangles and phage data with circles. For each host/phage group, %GC data is shown in a lighter shade than nRCA data. Data for hosts, for actinobacteriophage supercluster clusters (BI^*^, AM^*^, AU^*^, AW^*^, DJ^*^, CC^*^ and EL^*^) and for the *Faecalibacterium* and *Lactococcus* clusters studied here (Faec^*^ and Lacto^*^) are highlighted. Computations for *Faecalibacterium* hosts used available whole genome shotgun assemblies. All phage and host information is available in Table S4.

We also analyzed the codon usage bias (CUB) of phages with respect to their host (Figure 5), using the nRCA index (O’Neill, Or, and Erill 2013). In contrast to CAI, which is heavily influenced by mutational bias (Figure S6), the nRCA index explicitly corrects for base composition and hence primarily reflects bias linked to optimization for translational throughput. In each genus, as it is the case for %GC content, clusters within the actinobacteriophage supercluster display significantly lower average nRCA values than their hosts (4-30%, p<0.05), and significantly lower (p<0.05) average nRCA values than other phages infecting the same genera. In contrast, *Faecalibacterium* phage FP_oengus and *Lactococcus lactis* phages do not present significant differences in nRCA values with respect to their hosts or to other phages infecting them.

## Discussion

Bacteriophages will often infect several different hosts within the same bacterial genus, and this host range can vary widely among phages within a given genus (Erill and Caruso 2016; Caruso et al. 2019; Ross, Ward, and Hyman 2016). As a consequence, it has been postulated that the intragenera host–phage interaction network is nested, with generalist phages infecting multiple hosts and specialist phages infecting particularly susceptible strains (Flores et al. 2011). In contrast, relatively little is known about the ability of bacteriophages to infect across genera or broader taxonomic spans. Using plasmid-based transfer systems and multi-host isolation methods, phages capable of transcending genus boundaries have been selected (Ross, Ward, and Hyman 2016; Murooka, Takizawa, and Harada 1981), and effective transfer of virus-like particles via transduction has been documented across phyla (Chiura 1997). However, the occurrence in a natural setting of infections across distantly related bacterial groups has not been demonstrated. The recent availability of a significantly large amount of complete bacteriophage genomes infecting a wide variety of bacterial hosts provides an opportunity to explore the genetic relationship among bacteriophages infecting distantly related hosts, and to assess the possibility of such distant transfer events.

The identification of unexpected sequence similarity between orthologous protein sequences of phages infecting distantly related bacterial hosts within the Actinobacteria and the Firmicutes phyla led us to systematically explore their phylogenetic relationship. Both the gene content and terminase protein sequence phylogenies reported here (Figure 3 and Figure 4) indicate that actinobacteriophages infecting hosts from five different bacterial families (*Gordoniaceae, Mycobacteriaceae, Micrococcaceae, Microbacteriaceae* and *Nocardiaceae*) in two bacterial orders (Corynebacteriales and Micrococcales) are more closely related to each other than to any other sequenced phage infecting their respective hosts, forming a host-heterogeneous supercluster. Furthermore, phylogenetic analyses also reveal that these actinobacteriophages are closely related to phages infecting two different Firmicutes orders (Lactobacillales and Clostridiales) and that these, in turn, are more closely related to the actinobacteriophage supercluster than to other Firmicutes-infecting phages. This close evolutionary relationship is mostly driven by the conservation of two large genomic blocks involving replication and structural proteins (Figure 2), suggesting that these constitute the genomic backbone for this heterogeneous group of phages. This hypothesis is consistent with the observation of substantial variability in the intervening region between both blocks among the closely related BI cluster phages (Figure 1).

Analysis of genomic %GC content in this group of related Actinobacteria and Firmicutes phages reveals that their %GC content is systematically lower than that of their host genera and than that of similar phages infecting those genera. While the difference in %GC content between phages and their hosts is relatively small for *Lactococcus* and *Faecalibacterium* phages (~13%) it becomes much larger for Actinobacteria phages and hosts (20-30%). This trend is parallelled by codon usage bias, with Actinobacteria phages displaying significantly lower CUB than their hosts, and *Lactococcus* and *Faecalibacterium* phages exhibiting CUB values well-aligned with their hosts. This indicates that Actinobacteria phages lag behind in the process of ameliorating their %GC content and codon usage. In conjunction with the inferred phylogenies, the %GC and CUB analysis results posit two alternative scenarios for the emergence of this heterogeneous group of related phages. On the one hand, the ancestors of this group might have originated in a Gram-positive host, possibly related to *Lactococcus*, and spread first to high %GC Firmicutes (e.g. *Faecalibacterium*) before jumping to Actinobacteria hosts. On the other hand, these results may indicate that the selective pressure faced by phages to optimize their codon usage, and %GC content, to match the host’s in order to maximize translational throughput may be remarkably different for Actinobacteria- and Firmicutes-infecting phages. In bacteria, codon optimization for enhanced translational throughput is highly correlated with growth rate in laboratory settings, in which most Actinobacteria are known to grow rather slowly when compared to Firmicutes (Vieira-Silva and Rocha 2010; O’Neill, Or, and Erill 2013). Recent results indicate that this disparity in growth rates between Firmicutes and Actinobacteria extends to the wild, with Firmicutes often alternating dormant states with fast growing spurts and Actinobacteria seemingly replicating at lower, steadier rates (Brown et al. 2016; Gibson et al. 2018). This suggests that translational selection may be weaker in the Actinobacteria and their phages, resulting in lower rates of genome amelioration in Actinobacteria-infecting phage genomes, as reflected both in %GC content and codon usage bias profiles.

Recent analyses of genetic diversity in mycobacteriophages have put forward the notion that bacteriophages infecting mycobacteria do not constitute an isolated environment. Instead, rarefaction analyses suggest that the mycobacteriophage gene pool is constantly enriched by an influx of genetic material from external sources (Pope et al. 2015). The identification here of a group of related phages spanning multiple families within the Actinobacteria and encompassing also two Firmicutes orders suggests that, either through gradual evolution or host transfer, ancient phage lineages permeate phylum boundaries, thus contributing to the systematic enrichment of the gene pool available within the population of phages infecting any given genus. Lastly, it should be noted that the actinobacteriophage clusters identified here are not the only outliers in terms of %GC content and CUB divergence from their hosts, suggesting that further sequencing may enable the identification of other close evolutionary relationships between bacteriophages infecting distantly-related hosts.

## Author contributions

conceptualization, IE and SMC; methodology, IE and SMC; software, IE, MK, JLP.; validation, IE, SMC, MK, JLP and CM.; formal analysis, IE, MK; investigation, IE, SMC, CM, JLP, MK; resources, SMC, IE; data curation, IE, SMC, JLP, MK; writing—original draft preparation, IE; writing—review and editing, IE, SMC, CM, JLP, MK; visualization, IE, JLP, MK; supervision, IE and SMC; funding acquisition, SMC.

## Data accessibility

Data are available online: ZENODO repository DOI:<u>10.5281/zenodo.4645746</u>

## Supplementary material

Script and codes are available online: ZENODO repository DOI:<u>10.5281/zenodo.4645746</u>

## Acknowledgements

The authors wish to thank Ralph Murphy for his excellent technical support. The authors also wish to thank Graham F. Hatfull, Deborah Jacobs-Sera, Welkin H. Pope, Daniel R. Russell, Steven G. Cresawn and the Howard Hughes Medical Institute SEA-PHAGES program for their support. Version 3 of this preprint has been peer-reviewed and recommended by Peer Community In Genomics (https://doi.org/10.24072/pci.genomics.100003).

## Appendix

### Supporting Information Captions

**Figure S1** - Section of the multiple sequence alignment of the FP_oengus (AUV56548.1) gene product annotated as “putative RNA polymerase. Signature motifs shared by RDRPs and DNA-dependent RNA polymerases are highlighted in red. The signature metal-binding DxDxD motif is thought to be part of the primary catalytic loop for these RNA polymerases.

**Figure S2** - Consensus network inferred on SplitTree with the NeighborNet algorithm using a gene content distance. The branches corresponding to the actinobacteriophage supercluster and the *Lactococcus* and *Faecalibacterium* phage group are highlighted in red.

**Figure S3** - Phylogenetic tree from Neighbor Joining inference on the multiple sequence alignment of TerL sequences (Data S2).

**Figure S4** - Phylogenetic tree from minimal evolution inference on BLAST-derived inter-genomic distances. The branches corresponding to the actinobacteriophage supercluster and the *Lactococcus* and *Faecalibacterium* phage group are highlighted in red.

**Figure S5** - Detail of the reference viral proteomic tree generated by VipTree, highlighting the clustering of actinobacteriophage supercluster and the *Lactococcus* phages.

**Figure S6** - Average %GC and CAI of phage cluster genomes and of complete genomes from each cluster host genus. Cluster designations for actinobacteriophages are as assigned by PhagesDB. Host data is shown using triangles and phage data with circles. Data for hosts, for actinobacteriophage supercluster clusters (BI, AM, AU, AW, DJ, CC and EL) and for the *Faecalibacterium* and *Lactococcus* clusters studied here (Faec^*^ and Lacto^*^) are highlighted. Computations for *Faecalibacterium* hosts used available whole genome shotgun assemblies. All phage and host information is available in Table S4.

**Table S1** - Groups of orthologous proteins (phams) in the set of analyzed phage genomes.

**Table S2** - List of phage genomes analyzed in phylogenetic analyses.

**Table S3** - Average fraction of protein clusters (PCs) shared between members of the reported supercluster and versus other phages showing a significant fraction of shared protein clusters with them, as reported by Jang H, Bolduc B, Zablocki O, Kuhn JH, Roux S, Adriaenssens EM, *et al. Nat Biotechnol*. 2019;37: 632–639. doi:10.1038/s41587-019-0100-8.

**Table S4** - %GC content, nRCA and CAI values of phages and their hosts.

**Data S1** - FASTA-formatted files for host nRCA reference sets inferred with scnRCA.

**Data S2** - FASTA-formatted file with TermL sequences for the terminase tree.

**Data S3** - Nexus-formatted SplitTree file for the pham-based tree.

**Data S4** - Newick-formatted file for the TerL NJ tree.

