## Supplementary material for "Evidence for shared ancestry between Actinobacteria and Firmicutes bacteriophages": Figure S1

**Figure S1 –** Section of the multiple sequence alignment of the FP_oengus (AUV56548.1) gene product annotated as "putative RNA polymerase. Signature motifs shared by RDRPs and DNA-dependent RNA polymerases are highlighted in red. The signature metal-binding DxDxD motif is thought to be part of the primary catalytic loop for these RNA polymerases.

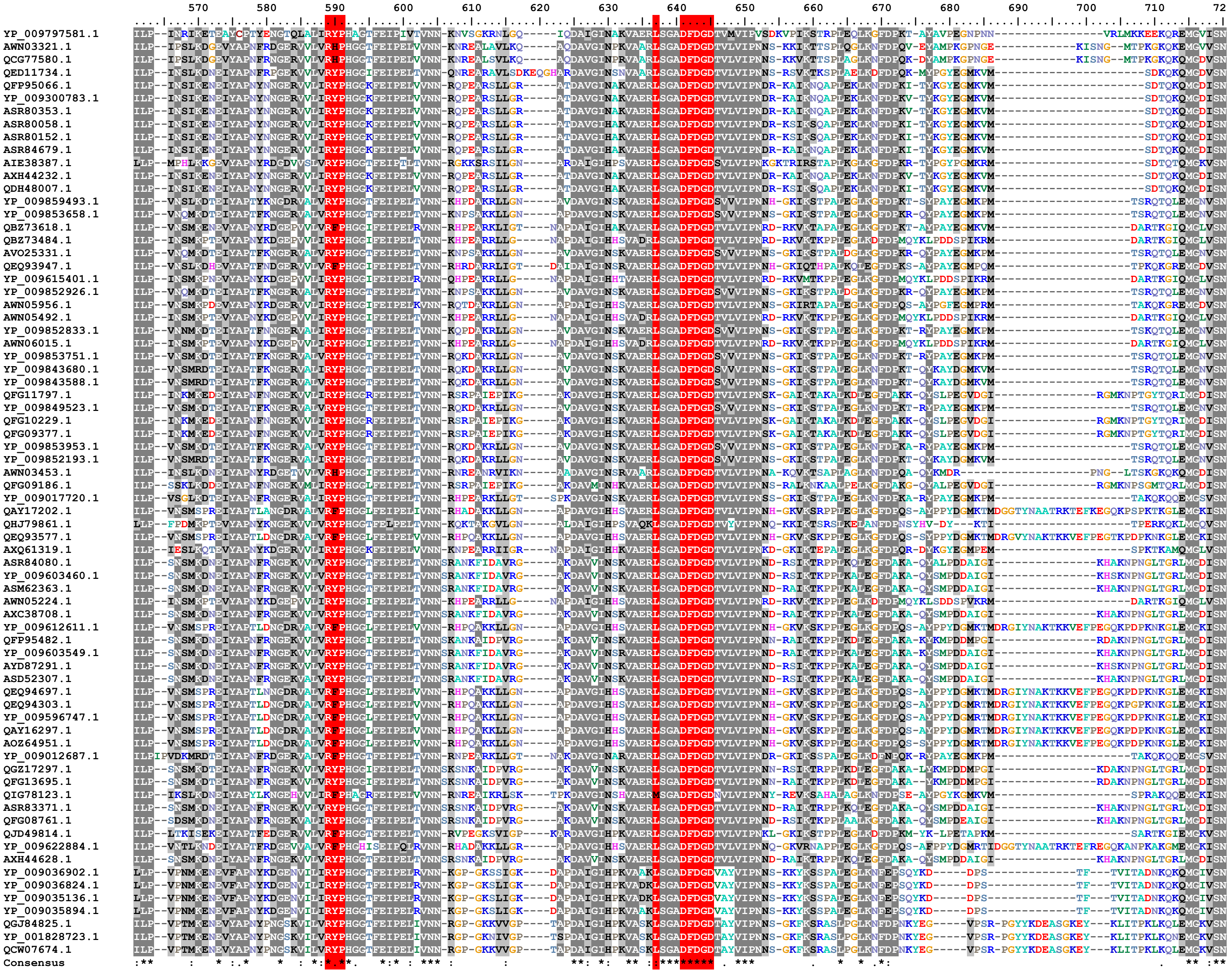

### References:


doi: 10.1038/s41564-017-0053-y

doi: 10.1186/1472-6807-3-1
