## Supplementary figures and images for "Evidence for shared ancestry between Actinobacteria and Firmicutes bacteriophages"

### Figure S2

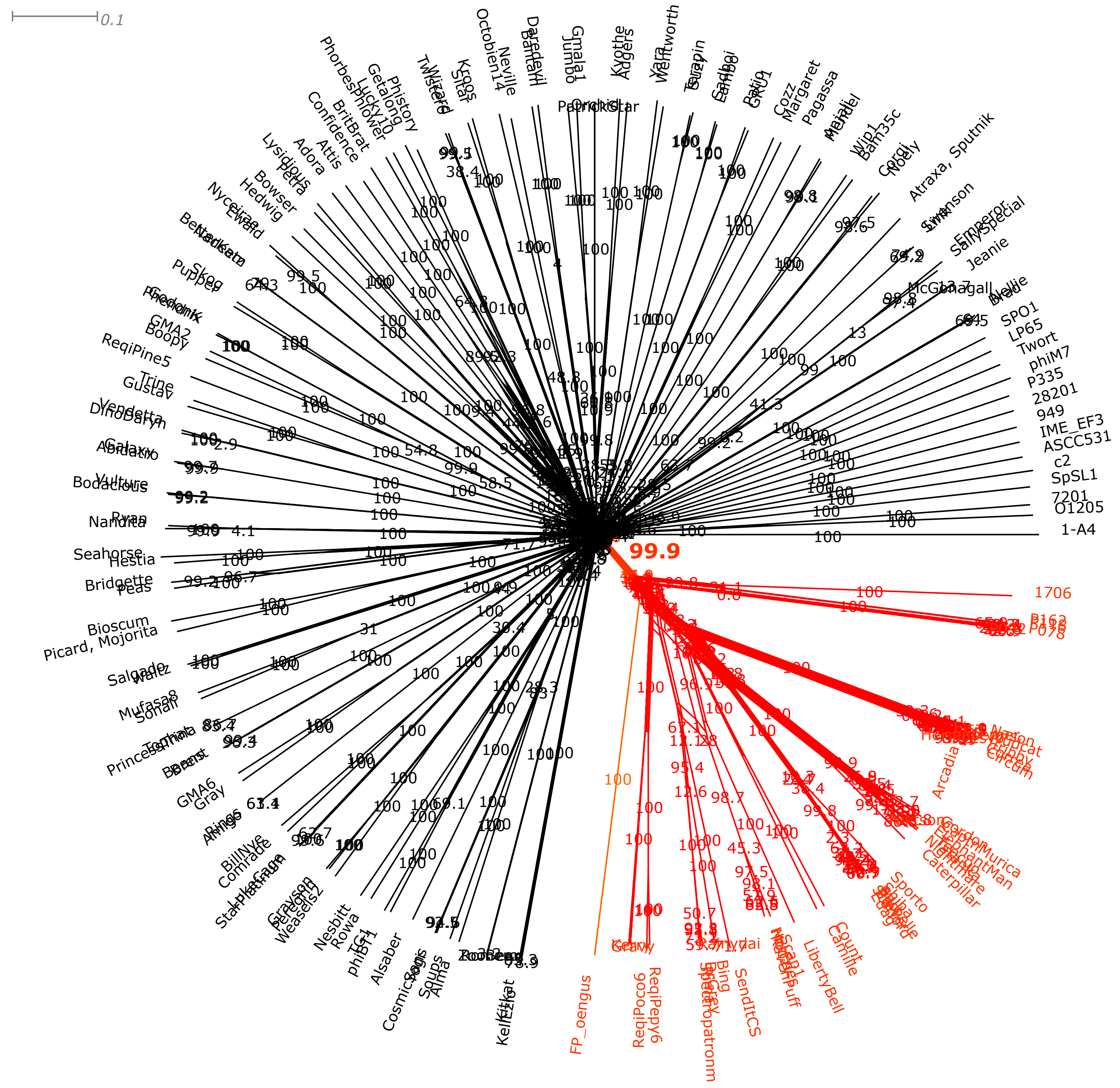

### Figure S3

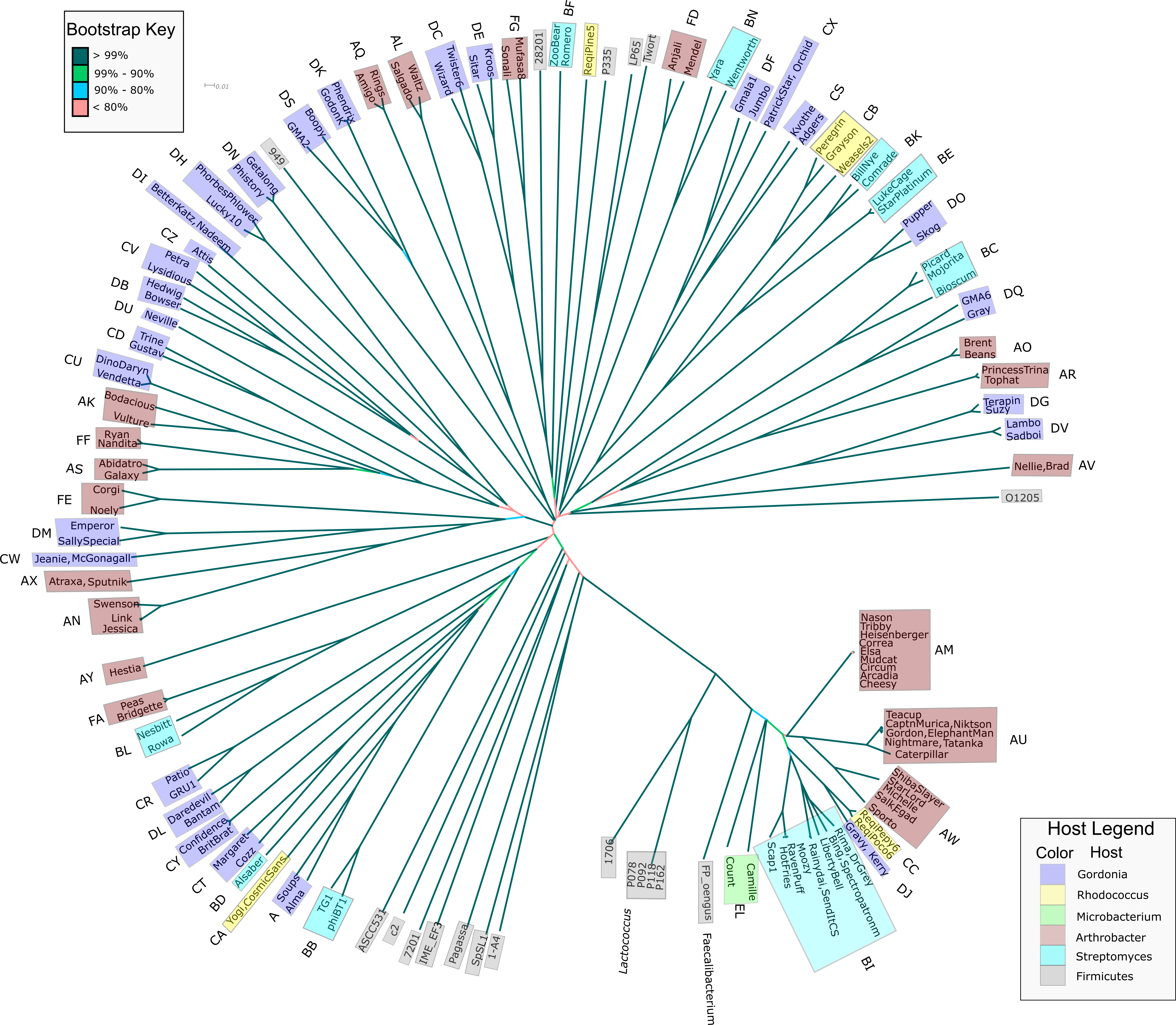

### Figure S4

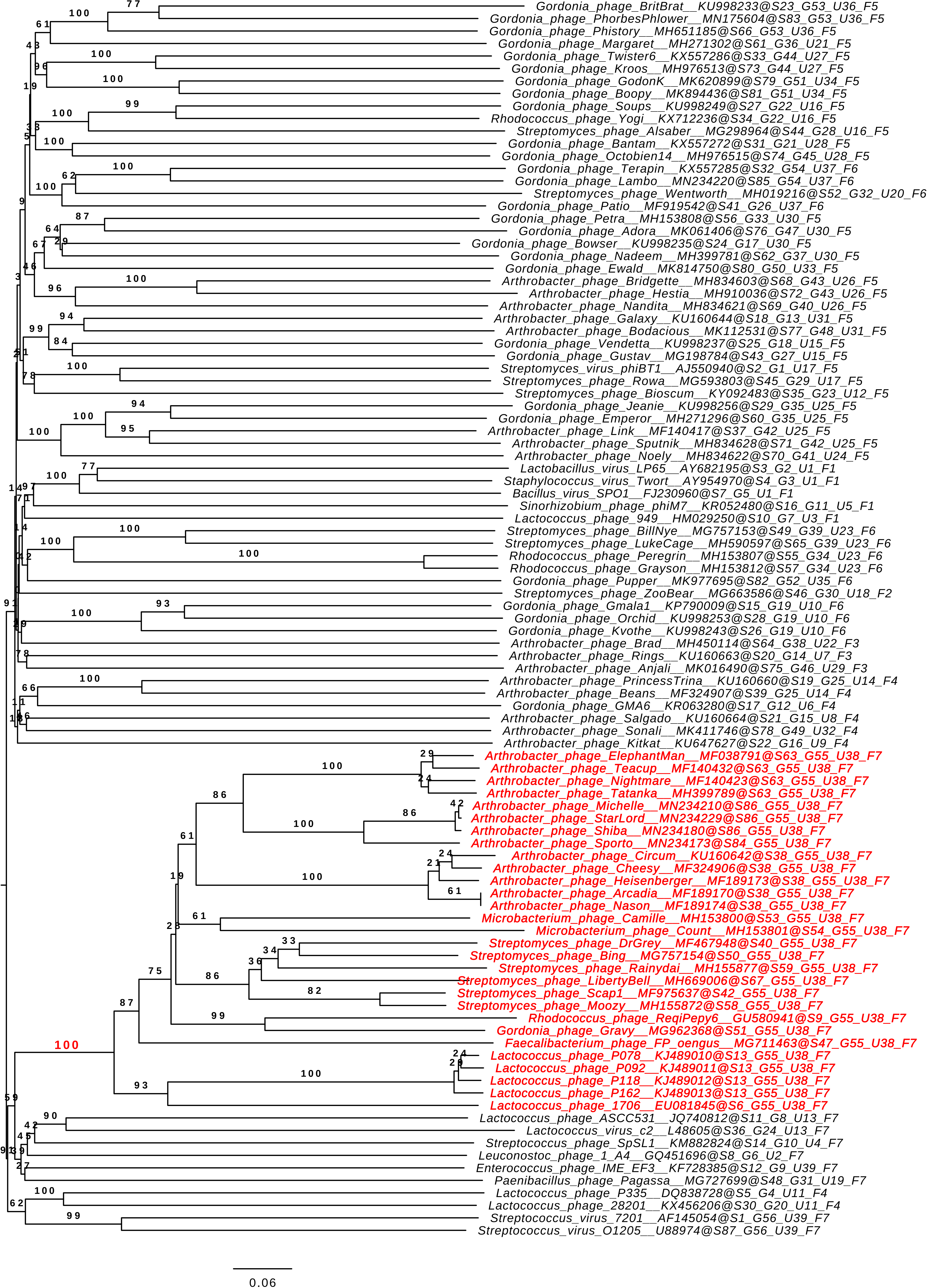

### Figure S5

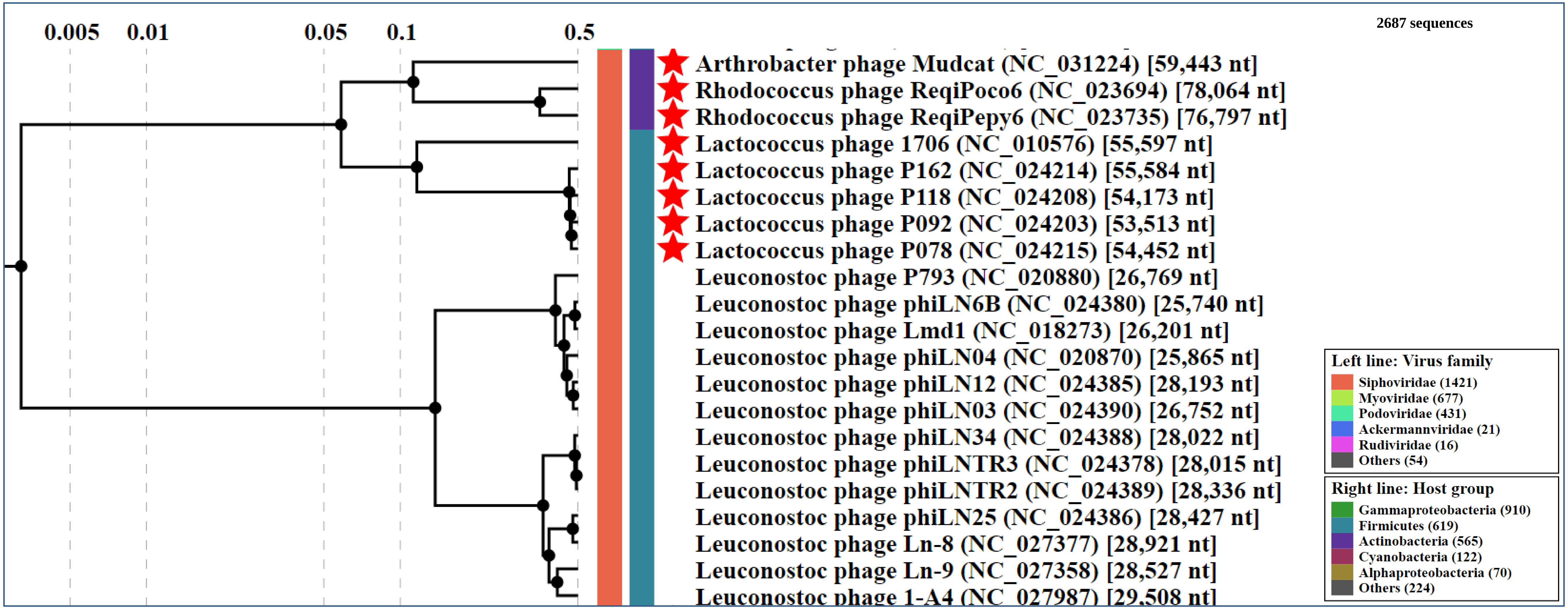

### Figure S6

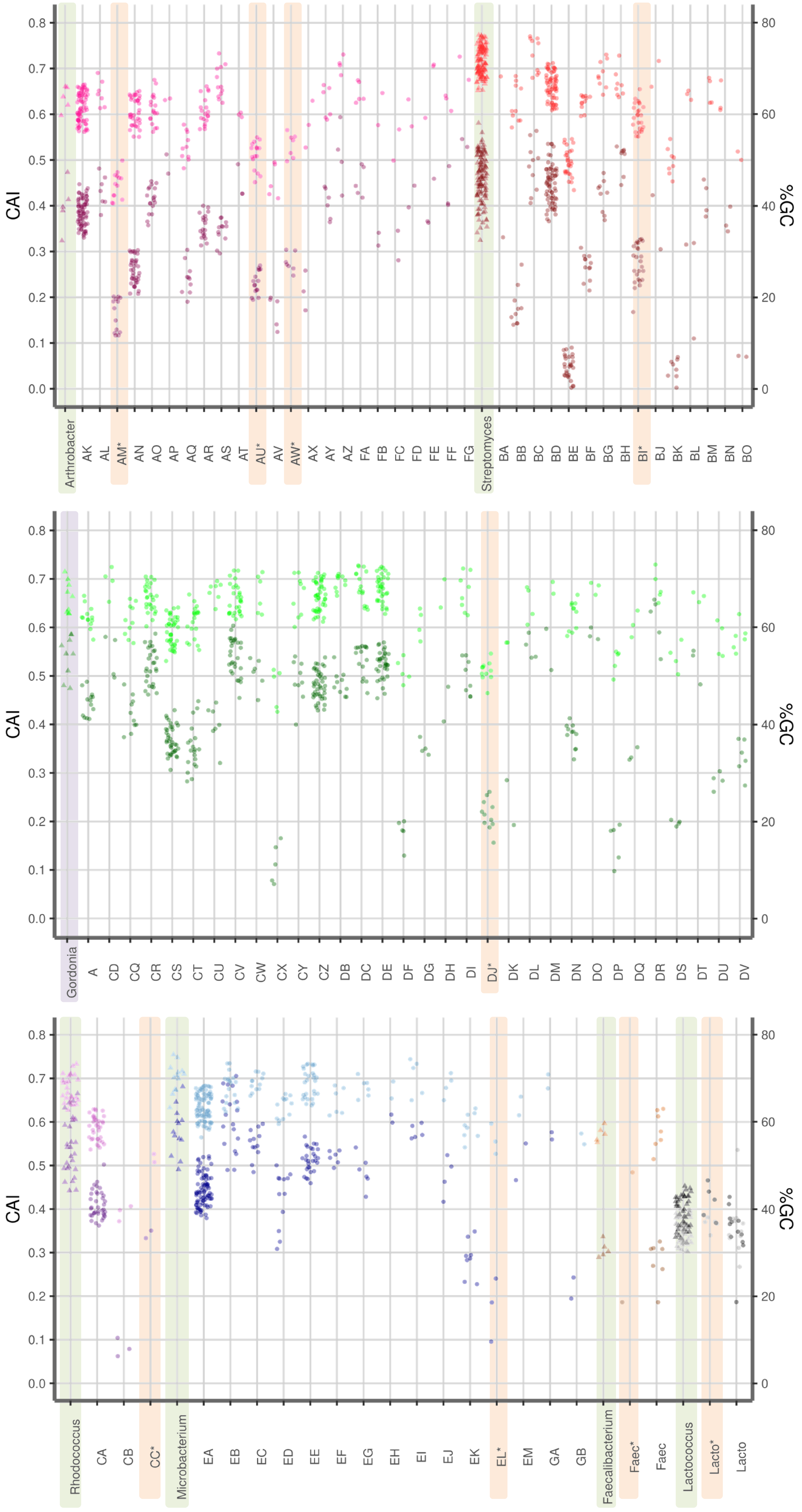
